# *K. pneumoniae* ST258 genomic variability and bacteriophage susceptibility

**DOI:** 10.1101/628339

**Authors:** Carola Venturini, Nouri Ben Zakour, Bethany Bowring, Sandra Morales, Robert Cole, Zsuzsanna Kovach, Steven Branston, Emma Kettle, Nicholas Thomson, Jonathan Iredell

**Affiliations:** Centre for Infectious Diseases and Microbiology, The Westmead Institute for Medical Research, The University of Sydney and Westmead Hospital, Sydney, Australia; AmpliPhi Australia Pty Ltd, Sydney, Australia; Westmead Research Hub Electron Microscope Core Facility, The Westmead Institute for Medical Research, The University of Sydney, Sydney, Australia; Parasites and Microbes, Wellcome Trust Sanger Centre, Cambridge, United Kingdom; The London School of Hygiene and Tropical Medicine, London, United Kingdom

**Author notes:** These authors contributed equally to this work (Carola Venturini, Nouri Ben Zakour). **Corresponding authors:** Carola Venturini, Centre for Infectious Diseases and Microbiology, The Westmead Institute for Medical Research, Westmead NSW 2145, Australia.; Jonathan R Iredell Centre for Infectious Diseases and Microbiology, The Westmead Institute for Medical Research, Westmead NSW 2145, Australia.

## Abstract

Multidrug resistant carbapenemase-producing *Klebsiella pneumoniae* capable of causing severe disease in humans is classified as an urgent threat by health agencies worldwide. Bacteriophages are being actively explored as potential therapeutics against these multidrug resistant pathogens. We are currently developing bacteriophage therapy against carbepenem-resistant *K. pneumoniae* belonging to the genetically diverse, globally disseminated clonal group CG258. In an effort to define a robust experimental approach for effective selection of lytic viruses for therapy, we have fully characterized the bacterial genomes of 18 target strains, tested them against novel lytic bacteriophages, and generated phage-susceptibility profiles. The genomes of *K. pneumoniae* isolates carrying *bla*_NDM_ and *bla*_KPC_ were sequenced and isolates belonging to CG258 were selected for susceptibility testing using a panel of lytic bacteriophages (n=65). The local *K. pneumoniae* CG258 population was dominated by isolates belonging to sequence type ST258 clade 1 (86%). The primary differences between ST258 genomes were variations in the capsular locus (*cps*) and in prophage content. We showed that CG258-specific lytic phages primarily target the capsule, and that successful infection is blocked in many, post-adsorption, by immunity conferred by existing prophages. Five bacteriophages specifically active against *K. pneumoniae* ST258 clade 1 (n=5) belonging to the Caudovirales order were selected for further characterization. Our findings show that effective control of *K. pneumoniae* CG258 with phage will require mixes of diverse lytic viruses targeting all relevant *cps* variants and allowing for variable prophage content. These insights will facilitate identification and selection of therapeutic phage candidates against this serious pathogen.

**Importance:** Bacteriophages are natural agents that exclusively and selectively kill bacteria and have the potential to be useful in the treatment of multidrug resistant infections. *K. pneumoniae* CG258 is a main agent of life-threatening sepsis that is often resistant to last-line antibiotics. Our work highlights some key requirements for developing bacteriophage preparations targeting this pathogen. By defining the genomic profile of our clinical *K. pneumoniae* CG258 population and matching it with bacteriophage susceptibility patterns, we found that bacteriophage ability to lyse each strain correlates well with *K. pneumoniae* CG258 structural subtypes (capsule variants). This indicates that preparation of bacteriophage therapeutics targeting this pathogen should aim at including phages against each bacterial capsular subtype. This necessitates a detailed understanding of the diversity of circulating isolates in different geographical areas in order to make rational therapeutic choices.

## Introduction

*Klebsiella pneumoniae* is an important ubiquitous Gram-negative species capable of causing disease in both humans and animals (1). The rise in recent decades of *K. pneumoniae* that are multi-drug resistant (MDR), including to last-line antibiotics such as carbapenems, has resulted in the classification of this species as an urgent threat to human health by health agencies worldwide and its recognition as an important antimicrobial resistance reservoir (2,3). Carbapenemase-producing (CP) MDR strains, carrying the *bla*_KPC_ and *bla*_NDM_ genes, can be asymptomatic residents of the human gut and a major cause of serious nosocomial infections worldwide associated with high morbidity and mortality (4-7).

The genetically diverse clonal group CG258, comprising sequence types ST258, ST11, ST512 and a few other SNP variants, is largely responsible for the global dissemination of MDR CP-*K. pneumoniae* (6,7). Population studies looking at the genomes of CG258 have shown that diversification within this clonal group is linked to a series of large-scale genomic rearrangements and an apparent high frequency of recombination, some of which result in switching or variation in the capsule polysaccharide-encoding (*cps*) locus (6-8). On the basis of this variation, the ST258 group can be subdivided into two separate lineages clade 1 and clade 2. ST258 clade 2 strains have been the main cause of disease outbreaks worldwide (7,9), while clade 1 uniquely predominated a recent Australian outbreak (9).

Alternative or adjuvant therapies to antibiotics against MDR pathogens are urgently needed (2). Naturally occurring lytic bacterial viruses (bacteriophages, phages) were recognised as effective therapeutic agents in the first decades of the 20th century, but were little valued by Western medicine during the antibiotic era (10). The rise in multidrug resistance, however, has renewed the interest in their potential for both decontamination and eradication of pathogens refractory to antibiotics. Lytic bacteriophages against problematic bacterial species can be readily isolated, but medical applicability is hindered by limited understanding of key issues such as optimal clinical protocols, penetration, and resistance, as well as disappointing clinical trials using phage that have been associated with inconsistent protocols and poor targeting (10-12). Bacteriophages capable of lysing *K. pneumoniae*, including MDR strains have been described (13-15), with complete genomes for more than 80 full double-stranded (ds) DNA phages available in NCBI databases to date. However, no effective therapeutic product has yet reached the bedside.

We are currently exploring bacteriophage therapy against extended-spectrum-β-lactamase (ESBL) producing *Enterobacteriaceae* isolated in Australia from humans with the aim of defining a robust experimental protocol for the rapid design of effective targeted phage preparations. Here, we present the bacteriophage susceptibility profiles of *K. pneumoniae* ST258 isolated in Australia and show their correlation with genomic variation within this clonal group. We also report the full genome sequence of five *K. pneumoniae* ST258-specific lytic phages (AmPh_EK29, AmPh_EK52, AmPh_EK80, JIPh_Kp122, JIPh_Kp127) isolated from wastewater in Australia.

## Methods

### Bacterial isolates

In this study, we have fully characterized the bacterial genomes of ESBL *K. pneumoniae* CG258 strains from Australia (n=18, with n=16 CP-*K. pneumoniae*) and tested the infectivity of novel bacteriophages (n=65) selected from our existing libraries or isolated *de novo* from local environmental sources. All MDR *K. pneumoniae* isolates containing the carbapenem resistance genes commonly associated with CG258, *bla*_NDM_ or *bla*_KPC_ (16), in our extensive clinical collection were selected as potential target isolates for this study (Table 1).

**Table 1.**
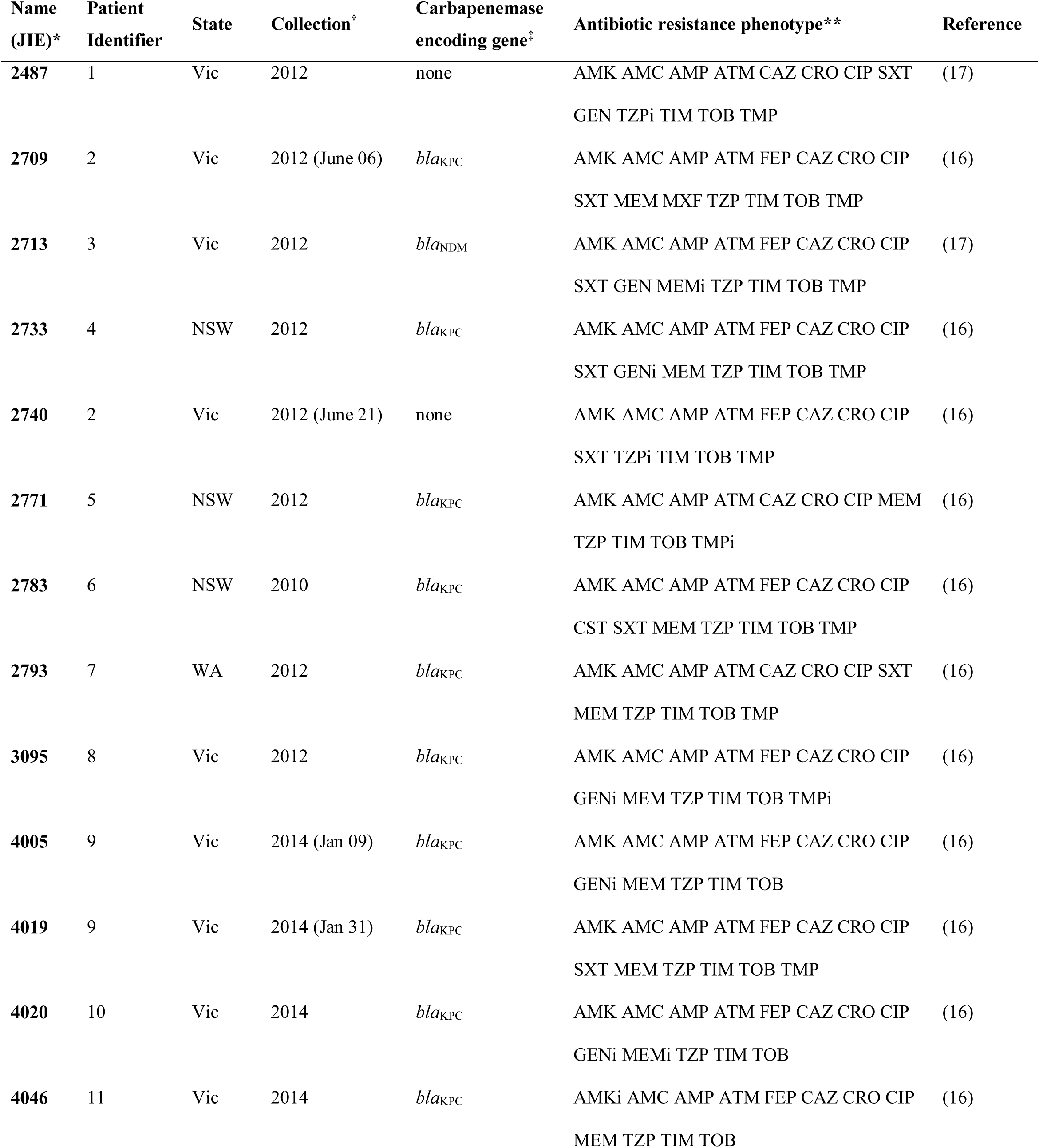
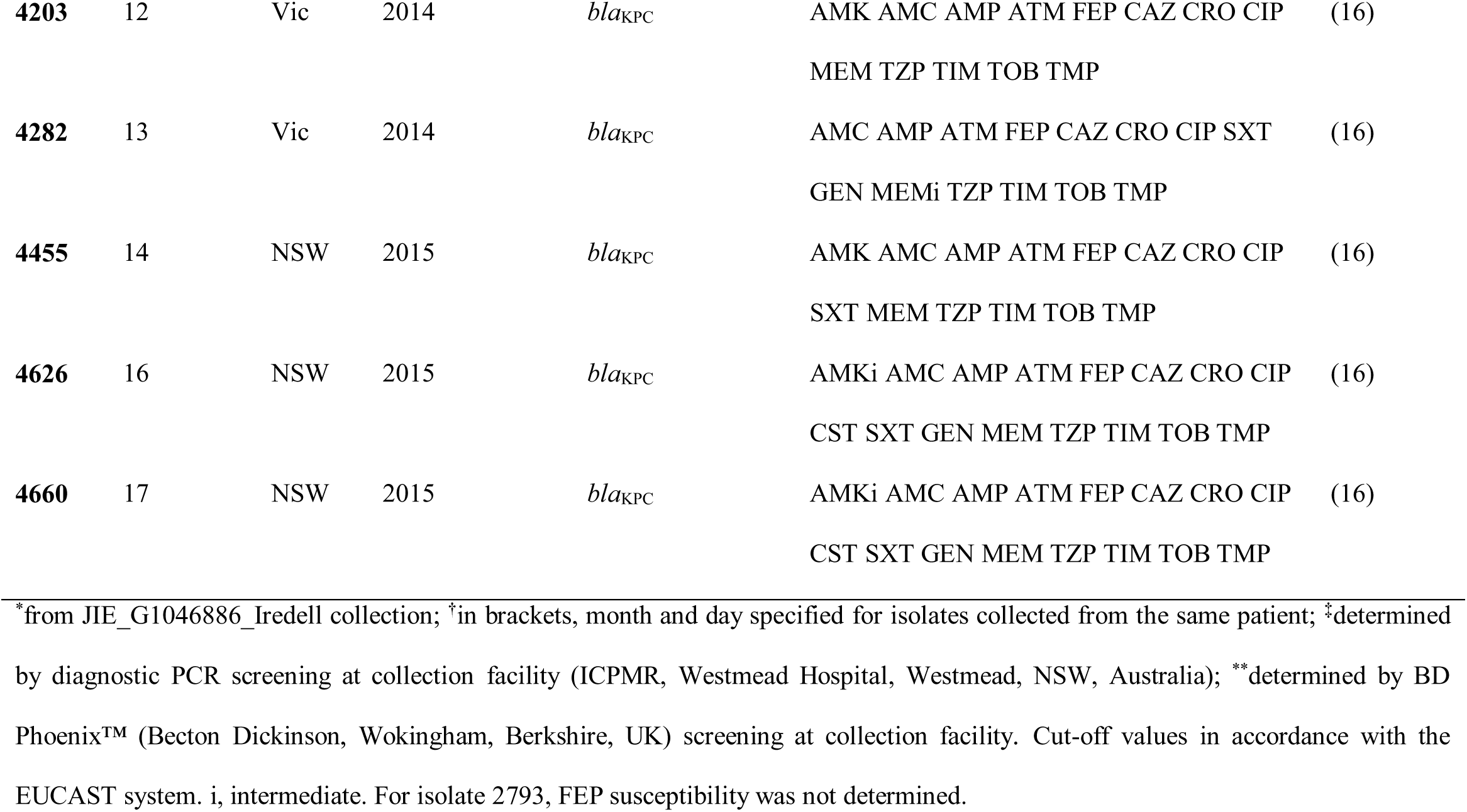
*K. pneumoniae* isolates characterized in this study.

| Name<br>(JIE)* | Patient<br>Identifier | State | Collection<sup>†</sup> | Carbapenemase<br>encoding gene<sup>‡</sup> | Antibiotic resistance phenotype** | Reference |
| --- | --- | --- | --- | --- | --- | --- |
| **2487** | 1 | Vic | 2012 | none | AMK AMC AMP ATM CAZ CRO CIP SXT<br>GEN TZPi TIM TOB TMP | (17) |
| **2709** | 2 | Vic | 2012 (June 06) | *bla<sub>KPC</sub>* | AMK AMC AMP ATM FEP CAZ CRO CIP<br>SXT MEM MXF TZP TIM TOB TMP | (16) |
| **2713** | 3 | Vic | 2012 | *bla<sub>NDM</sub>* | AMK AMC AMP ATM FEP CAZ CRO CIP<br>SXT GEN MEMi TZP TIM TOB TMP | (17) |
| **2733** | 4 | NSW | 2012 | *bla<sub>KPC</sub>* | AMK AMC AMP ATM FEP CAZ CRO CIP<br>SXT GENi MEM TZP TIM TOB TMP | (16) |
| **2740** | 2 | Vic | 2012 (June 21) | none | AMK AMC AMP ATM FEP CAZ CRO CIP<br>SXT TZPi TIM TOB TMP | (16) |
| **2771** | 5 | NSW | 2012 | *bla<sub>KPC</sub>* | AMK AMC AMP ATM CAZ CRO CIP MEM<br>TZP TIM TOB TMPi | (16) |
| **2783** | 6 | NSW | 2010 | *bla<sub>KPC</sub>* | AMK AMC AMP ATM FEP CAZ CRO CIP<br>CST SXT MEM TZP TIM TOB TMP | (16) |
| **2793** | 7 | WA | 2012 | *bla<sub>KPC</sub>* | AMK AMC AMP ATM CAZ CRO CIP SXT<br>MEM TZP TIM TOB TMP | (16) |
| **3095** | 8 | Vic | 2012 | *bla<sub>KPC</sub>* | AMK AMC AMP ATM FEP CAZ CRO CIP<br>GENi MEM TZP TIM TOB TMPi | (16) |
| **4005** | 9 | Vic | 2014 (Jan 09) | *bla<sub>KPC</sub>* | AMK AMC AMP ATM FEP CAZ CRO CIP<br>GENi MEM TZP TIM TOB | (16) |
| **4019** | 9 | Vic | 2014 (Jan 31) | *bla<sub>KPC</sub>* | AMK AMC AMP ATM FEP CAZ CRO CIP<br>SXT MEM TZP TIM TOB TMP | (16) |
| **4020** | 10 | Vic | 2014 | *bla<sub>KPC</sub>* | AMK AMC AMP ATM FEP CAZ CRO CIP<br>GENi MEMi TZP TIM TOB | (16) |
| **4046** | 11 | Vic | 2014 | *bla<sub>KPC</sub>* | AMKi AMC AMP ATM FEP CAZ CRO CIP<br>MEM TZP TIM TOB | (16) |

Table 1 continued
| Name<br>(JIE)* | Patient<br>Identifier | State | Collection<sup>†</sup> | Carbapenemase<br>encoding gene<sup>‡</sup> | Antibiotic resistance phenotype** | Reference |
| --- | --- | --- | --- | --- | --- | --- |
| 4203 | 12 | Vic | 2014 | *bla<sub>KPC</sub>* | AMK AMC AMP ATM FEP CAZ CRO CIP<br>MEM TZP TIM TOB TMP | (16) |
| 4282 | 13 | Vic | 2014 | *bla<sub>KPC</sub>* | AMC AMP ATM FEP CAZ CRO CIP SXT<br>GEN MEMi TZP TIM TOB TMP | (16) |
| 4455 | 14 | NSW | 2015 | *bla<sub>KPC</sub>* | AMK AMC AMP ATM FEP CAZ CRO CIP<br>SXT MEM TZP TIM TOB TMP | (16) |
| 4626 | 16 | NSW | 2015 | *bla<sub>KPC</sub>* | AMKi AMC AMP ATM FEP CAZ CRO CIP<br>CST SXT GEN MEM TZP TIM TOB TMP | (16) |
| 4660 | 17 | NSW | 2015 | *bla<sub>KPC</sub>* | AMKi AMC AMP ATM FEP CAZ CRO CIP<br>CST SXT GEN MEM TZP TIM TOB TMP | (16) |

\*from JIE\_G1046886\_Iredell collection; <sup>†</sup>in brackets, month and day specified for isolates collected from the same patient; <sup>‡</sup>determined by diagnostic PCR screening at collection facility (ICPMR, Westmead Hospital, Westmead, NSW, Australia); \*\*determined by BD Phoenix™ (Becton Dickinson, Wokingham, Berkshire, UK) screening at collection facility. Cut-off values in accordance with the EUCAST system. i, intermediate. For isolate 2793, FEP susceptibility was not determined.

### CP-*K. pneumoniae* phenotypes

#### Biofilm production

Biofilm formation by growing bacteria in polypropylene microtitre plates was estimated by crystal violet staining of adherent cells, following the protocol of O’Toole and Kolter (18) with minor modifications. Briefly, overnight bacterial cultures in lysogeny broth (LB; Oxoid, Basingstoke, UK) adjusted to OD_600_ 0.4 were added (0.1 mL) to microtiter plate (Corning Life Sciences, Corning, NY, USA) wells and grown overnight in a static incubator at 37°C. Wells were carefully washed twice with RO water before addition of 0.1% crystal violet (Sigma-Aldrich, MO, USA) (225 μL) and incubation at room temperature. Plates were gently washed four times with RO water and dried at room temperature for at least 2 h. For quantitation, 200 µL of 100% ethanol was added to each well and left for 10-15 min. An aliquot (125 μL) of the solubilized solution was then transferred to a new flat bottom microtiter dish and absorbance at 540 nm was measured in a Spectromax Vmax microplate reader (Biomolecular Devices, San Jose, CA, USA). Negative (LB only) and positive (ATCC 27853 *P. aeruginosa*, a strong biofilm producer) controls were included on all plates. Experiments were performed in triplicate.

#### Polysaccharide capsule production

Total capsule production in *K. pneumoniae* CG258 was quantified according to previously described methods (19,20). Briefly, overnight bacterial cultures in Mueller-Hinton broth (Oxoid, Basingstoke, UK) were mixed with 1% Zwittergent 3-14 detergent (Millipore, Billerica, MA, USA) in 100 mM citric acid (pH 2.0) and incubated for 30 min at 50°C with occasional mixing. After pelleting the bacteria, 300 µL of supernatant was mixed with absolute ethanol to a final concentration of 80% and left on ice for 30 min to allow for capsule precipitation. After centrifugation, the precipitates were allowed to dry and then resuspended in 100 µL of DNase-free water (Lonza, Rockland, ME, USA) and kept at 4°C overnight. Capsule quantitation was assayed by measuring uronic acid content on ethanol-precipitated culture supernatants by addition of 1.2 mL of 12.5 mM tetraborate (Sigma-Aldrich, St. Louis, MO, USA) in concentrated sulphuric acid (Sigma-Aldrich, St. Louis, MO, USA), and detection (absorbance at 520 nm) using 0.15% m-hydroxydiphenyl (Sigma-Aldrich, St. Louis, MO, USA) in 0.5% NaOH (Amresco, Solon, OH, USA). Sodium hydroxide added to the tetraborate/sulphuric acid solution was used as the baseline for quantification. Capsule quantification was performed in triplicate for each bacterial strain.

#### Lipopolysaccharide profiles

Lipopolysaccharide (LPS) profiles of *K. pneumoniae* CG258 strains were analysed using sodium dodecyl sulphate polyacrylamide gel electrophoresis (SDS-PAGE) subsequent to proteinase K digestion of the proteins in whole cell lysates (21). Overnight bacterial cultures (100 µl) were pelleted and washed twice in saline. The final pellet was resuspended in 35 µl sterile saline. Pellets were treated with 20 µl 4× SDS reducing buffer (0.0625 M Tris-HCl, pH 8.8, 10% glycerol, 2% SDS, 5% 2-b-mercaptoethanol, 0.0125% bromophenol blue) at 100°C for 10 min. Proteins were digested by addition of 15 µl 20 mg/ml proteinase K and incubation at 60°C for 1 h. Sample preparations were run on a 15% separating gel with a 4% stacking gel under reducing conditions using Tris-glycine running buffer (mini-PROTEAN system, Bio-Rad Laboratories, Hercules, CA, USA). Silver staining using sodium thiosulphate sensitisation and silver nitrate was performed according to established methods (22).

### Sequencing and analysis of bacterial genomes

The genomes of selected MDR *K. pneumoniae* isolates were sequenced by Illumina NextSeq (paired-end; 2 × 150 bp). Bacterial DNA extraction was performed using the DNeasy Blood and Tissue DNA isolation kit (Qiagen, Hilden, Germany) to obtain high purity (OD_260/280_ 1.8-2.0; OD_260/230_ 1.8) preparations for sequencing. DNA libraries for whole genome sequencing (WGS) were prepared using the Nextera XT kit and sequencing was performed at the Australian Genome Research Facility (AGRF, Melbourne, Australia). *De novo* assembly of sequencing reads and simulated reads of NCBI reference genomes were performed as previously described (23), using our WGS analysis workflow based on publicly available tools including SPAdes 3.9.0 (24); Nullarbor 1.20 (25); Kleborate 0.2.0 (26) to confirm identity (*in silico* MLST), virulence and antibiotic resistance genotypes. A maximum-likelihood recombination-free phylogenetic tree was computed using RAxML 8.2.4 (27) and Gubbins 2.2.0 (28), using a reference-based core genome alignment as an input. The publicly available genome sequences of five representative CG258 strains were also added for comparative purposes: AUSMDU00008079, used as mapping reference (CP022691); HS11286 (CP003200); NJST258-1 (CP006923); Kb140 (AQROD00000000); and VA360 (ANGI00000000). The pangenome was determined using Roary version 3.11.0 (29) and used to classify regions of differences across the strain dataset, based on their contiguity and functional categories of the genes encoded. Kleborate 0.2.0 (26) was used to perform capsule typing, O antigen (LPS) serotyping and siderophore typing. Plasmid replicon identification and typing was performed using PlasmidFinder and pMLST implemented in BAP (30). Prophage-associated contigs were annotated using PHASTER (31). Further *cps* locus comparative analysis was performed using EasyFig (32) and Geneious v9.1 (https://www.geneious.com). Gap closure between separate contigs in the capsular locus was achieved by PCR amplification and Sanger sequencing of purified linkage amplicons (AGRF, Melbourne, Australia).

#### bla_KPC_ genomic context

To confirm plasmid content and genomic context of the *bla*_KPC_ gene in target ST258 (n=16) isolates, we performed Pulse Field Gel Electrophoresis (PFGE) on S1 nuclease (Promega, Madison, WI, USA) digested DNA, as before (33,34), and Southern hybridization with *bla*_KPC_ and *rep* IncFII_K_ DIG-labelled probes (16) prepared using published primers (35,36) and the PCR DIG Probe Synthesis Kit (Roche, Mannheim, Germany) following manufacturer’s instructions. Images were obtained on a ChemiDoc™ MP System (Bio-Rad Laboratories, Richmond, CA, USA).

### *De novo* isolation of CP-*K. pneumoniae* specific bacteriophages

Bacteriophages against target *K. pneumoniae* CG258 were isolated from sewage and wastewater samples collected in the Greater Sydney District (Sydney, NSW, Australia). Specimens were clarified by centrifugation and filtration through a 0.22 µm filter. Aliquots of environmental filtrates were then incubated with target *K. pneumoniae* isolates overnight. Bacteriophages were selected from single plaques in double-layer agar assays and purified through three rounds of plating (37). High-titre stocks were prepared by propagating phage over several double-layer plates washed in SM buffer (50 mM Tris-HCl, 8 mM MgSO_4_, 100 mM NaCl, pH 7.4) and filtered through a 0.22 µm filter. The concentration of phage-forming units (PFU) per ml was determined by spotting 10 µl of ten-fold serial dilutions onto a double-layer of the target bacteria (37). High-titre (≥ 10^9^ PFU/mL) phage stocks were stored at 4°C.

### Bacteriophage host range

The identified CG258 *K. pneumoniae* strains were tested against phages (n=65) selected from our extensive library or isolated *de novo* against one of the target isolates. Bacteriophage host range was determined by measuring the efficiency of plating (EOP) for each phage-bacteria combination. Ten-fold serial dilutions (10 µl) were spotted onto a double-layer of the target bacteria and compared to the original isolation host (37). *Escherichia coli, Enterococcus faecium, Pseudomonas aeruginosa, Staphylococcus aureus* and *Staphylococcus pseudintermedius* were used as cross-species controls. In order to further test bacteriophage specificity, we also blind tested the host ranges of three phages with unique specificities (AmPh_EK52, AmPh_EK38 and JIPh_Kp122) on a set of CP-*K. pneumoniae* isolates from Europe (n=48) to determine their predictive diagnostic value linked to *K. pneumoniae* sequence or capsule type. Bacteriophage activity against each strain was scored as: 1. ‘full activity’ for presence of clear plaques at highest dilution; 2. ‘poor activity’ for presence of turbid plaques, or isolated bacterial colonies within clearings, or EOP three or more log_10_ lower than that of the original host; 3. ‘partial activity’ for evidence of clearing in bacterial lawn, but absence of distinct plaques; 4. ‘negative’ for very faint, difficult to observe, clearing or absence of any visible plaques or clearing zones in the bacterial lawn.

### Bacteriophage characterization

#### Genome sequencing

Bacteriophages (n=5) that specifically lysed *K. pneumoniae* ST258 clade 1 with different host range profiles were selected for further characterization as potential therapeutic candidates. Bacteriophage DNA was extracted using the Wizard DNA Clean-Up System (Promega, Madison, WI, USA) and used for whole genome sequencing (WGS) (Nextera XT kit; Illumina NextSeq, paired-end, 2 × 150 bp). Briefly, total DNA concentration was quantified using Quant-it PicoGreen dsDNA Assay Kit (Invitrogen, Carlsbad, CA, USA) and 1 ng/µl of DNA was used to prepare DNA libraries using the Nextera XT Library Preparation Kit and Nextera XT v2 Indexes (Illumina, San Diego, CA, USA). Multiplexed libraries were sequenced using paired end 150 bp chemistry on the NextSeq 500 NCS v2.0 (Illumina). Error rates were calculated using PhiX Sequencing Control v3 for each run. De-multiplexing and FastQC generation was performed with default settings using BaseSpace (Illumina). Bacteriophage genomes were assembled using our in-house genomic pipeline and annotated using RAST-tk (38). The absence of lysogeny modules, virulence and resistance determinants was determined using our WGS analysis workflow (as for bacterial genomes) and PHASTER (31). Genome comparisons with best database (GenBank, NCBI) matches were obtained using EasyFig (32). PFGE of intact viral particles was performed to confirm relative size (Chef Mapper System, Bio-Rad Laboratories, Hercules, CA, USA).

#### Imaging of bacteriophages

Bacteriophage preparations were dialysed against 0.1 M ammonium acetate in dialysis cassettes with a 10,000 membrane molecular weight cut-off (Pierce Biotechnology, Rockford, IL, USA), negatively stained with 2% uranyl acetate and visualised using transmission electron microscopy (TEM) (37). TEM was conducted at the Westmead Electron Microscopy Facility (Westmead, Australia) on a Philips CM120 BioTWIN transmission electron microscope at 100kV. Images were recorded with a SIS Morada digital camera using iTEM software (Olympus Soft Imaging Solutions GmbH, Munster, Germany). Bacteriophage morphology and related taxonomic assignment were confirmed following the guidelines set by the International Committee on Taxonomy of Viruses (http://www.ictvonline.org/; 39).

#### Phage stability

Phage stability in SM buffer was determined by measuring the EOP after incubation at different temperatures (21°C for 24 h, 21°C for 7 days, 37°C for 24 h, 4°C for >1 month, and 4°C for >1 month with chloroform) and at different pH levels (pH 3, 6, 7, and 8; 4 h at room temperature). Rate of adsorption and one-step growth curves (latent period and burst size) were calculated according to established protocols (37).

## Results

### CP-*K. pneumoniae* population

From a total of 21 isolates sequenced, 18 ESBL *K. pneumoniae* belonged to CG258 (S1 Table). This set contained two ST11 representatives, one ST512 (single locus variant of ST258 clade 2) (7), one ST1199 (single SNP variant of ST258 clade 1) (40), and 14 ST258 strains, segregated further into two distinct lineages, known as clade 1 (n=13) and clade 2 (n=2) and distinguished by approx. 40 non-recombinant core SNPs (Fig 1). Strains belonging to clade 1 dominated this population (86%), reflecting the epidemiology of a recent Australian outbreak (9). Inspection of the most common *K. pneumoniae* virulence-associated mobile genetic element, ICE*Kp*, revealed that all strains belonging to ST11 carried an ICE*Kp3* associated with a yersiniabactin operon *ybt* 9, while all ST258 and the ST1199 strain carried an ICE*Kp2* associated with a yersiniabactin operon *ybt* 13, a combination usually only found in clade 1 isolates (Fig. 1) (clade 2 isolates most commonly being associated with ICE*Kp10* with *ybt* 17) (41).

**Figure 1.**
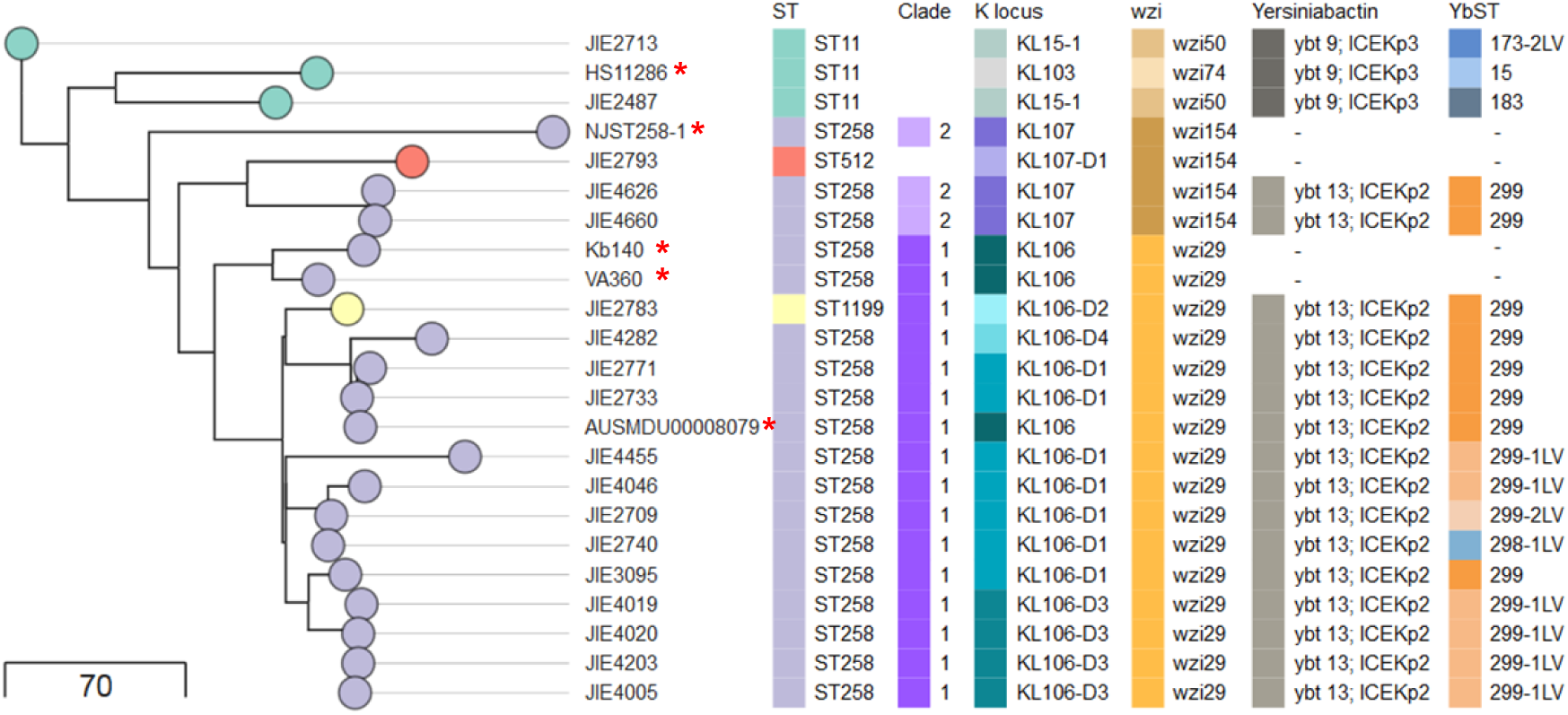
*K. pneumoniae* CG258 phylogeny. Recombination-free maximum-likelihood phylogenetic tree of 18 *K. pneumoniae* CG258 isolates and five publically available reference genomes (**\***). Metadata include MLST; CG258 clade; *cps* type (KL and *wzi*); Yersiniabactin, ICE*Kp* and ST (YbST). Tree scale corresponds to number of substitutions.

### CP-*K. pneumoniae* CG258 bacteriophage susceptibility

Screening for bacteriophage susceptibility identified at least one phage with strong activity against each strain of CP-*K. pneumoniae* CG258, with potential therapeutic value (Fig. 2). A correlation pattern between specific regions of differences (RODs) and bacteriophage susceptibility was evident in our dataset (Fig.2). Phage host range profiles largely grouped according to capsular types, KL15-1 (ST11), KL106-D1 (ST258 clade 1) and KL107 (ST258 clade 2 and ST512) each presenting unique patterns of phage susceptibility with few examples of cross-reactivity. Phage AmPh_EK29 was able to lyse both a subset of clade 1 isolates (KL106-D1 and D2) and JIE2793 (ST512; KL107), while phages JIPh_Kp122 and JIPh_Kp127 had preferential activity against few specific clade 1 isolates (Fig. 2).

**Figure 2.**
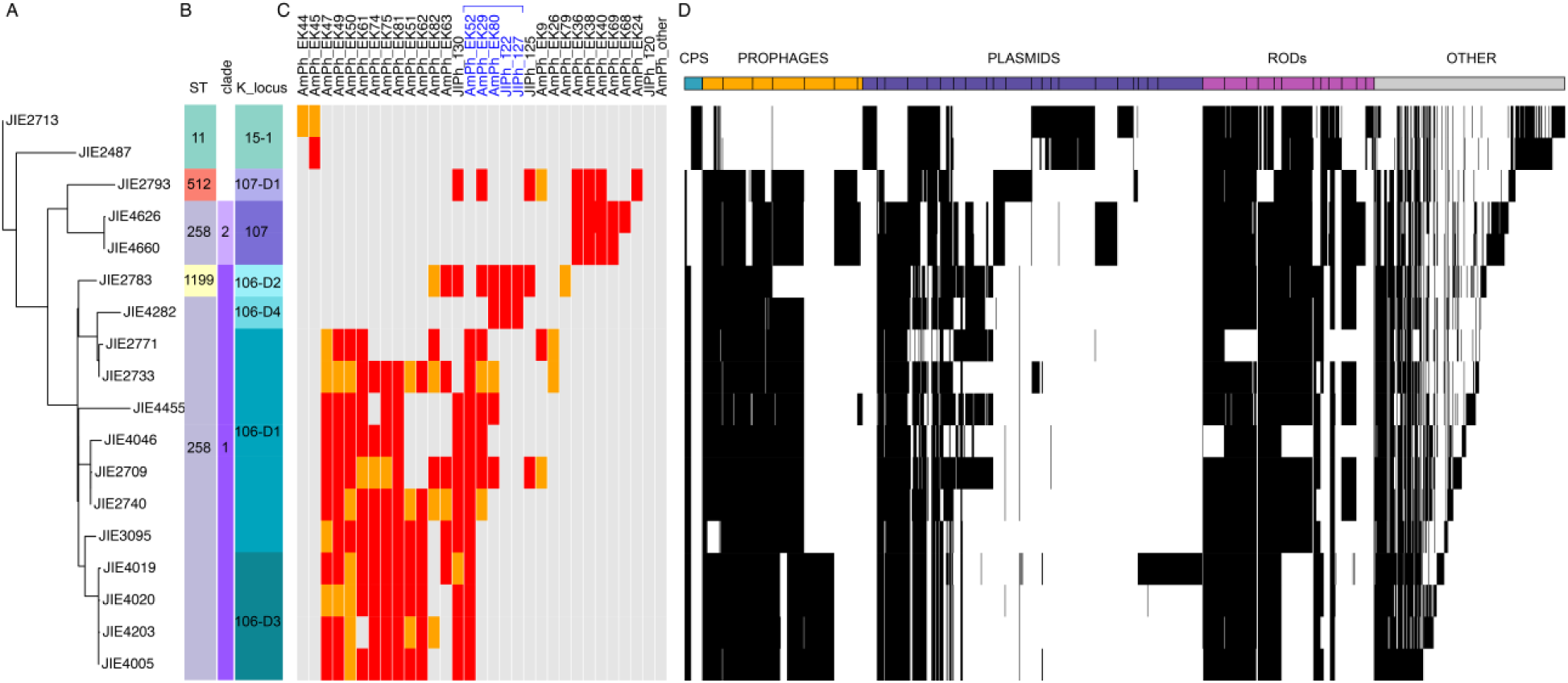
Phage susceptibility profiles and regions of differences. **A)** *K. pneumoniae* CG258 isolates phylogeny; **B)** metadata including ST, CG258 clade and K locus type; **C)** phage susceptibility profiles colour-coded based on therapeutic application potential as follows; grey, unsuitable for further testing (no lysis); orange, poor lytic activity; red, best candidates for further characterization (high lytic activity). Five phages (highlighted in blue) were selected for full characterization; **D)** regions of difference profiles identified using Roary (black, present). Genes corresponding to the accessory genome (n=1683) were reordered according to their synteny where possible, and classified according to functional categories into regions of difference as follows: CPS, capsule associated-genes; prophage-related regions; plasmid-related regions; other regions of differences (RODs) such as ICE elements; and other genes present in variable regions <10 consecutive genes. Details of regions are listed in Table S2.

Within the clade 1 set, this capsule-specificity of the phage tested was further confirmed by the resistance to lysis by the capsular variant isolates JIE2783 and JIE4282 (see below) which were resistant to the majority of the bacteriophages capable of lysing other clade 1 strains. *K. pneumoniae* phages AmPh_EK52 and JIPh_130 were effective in clearing most CP-*K. pneumoniae* ST258 clade 1 specifically, except for the ones carrying capsular variants. There was no cross-activity of any of these clade-specific phages against any of the control species tested (*E. coli, E. faecium, P. aeruginosa, S. aureus* or *S. pseudintermedius*).

### ESBL *K. pneumoniae* CG258 variable accessory genome

A comprehensive analysis of the pan-genome using Roary showed that the CG258 accessory genome was composed of 1,682 accessory genes, which could be further classified into discrete RODs (regions of difference) based on their function and contiguity as follows: 33 capsule-associated genes, 334 phage-related genes, 622 plasmid-related genes, 328 RODs-associated genes, and 365 other genes that could not be assigned unambiguously to the aforementioned categories (ROD were numbered and are summarised in S2 Table). The presence-absence profile of these RODs appeared to reflect the ST and/or clade of the isolates, in particular for prophage- and plasmid-related regions (Fig. 2ABD; S2 Table). Notwithstanding this, there was a substantial degree of intra-clade variation within the ST258 clade 1 isolates in the RODs corresponding to the *cps* polysaccharide capsule encoding locus, plasmid content and other RODs (Fig. 2ABD; S2 Table).

#### Polysaccharide capsule synthesis locus

Bioinformatic analysis of the genomic region between the *galF* and *ugd* genes revealed distinct capsular types within the same ST, with KL15 and KL103 found in ST11 strains, and KL106 and KL107 found in ST258 clade 1 and 2, respectively. The *cps* locus is a well-known recombination hot-spot in *K. pneumoniae* and further inspection of our clade 1 isolates revealed the presence of three variants of the previously described KL106-D1 arrangement (40,42) (Fig. 2; Fig. 3a). Variants were due to insertion of IS*Kpn26* (IS5 family) in two different locations within the *cps* locus: (i) within the *wcaJ* gene (KL106-D2 in JIE2783 and KL106-D4 in JIE4282) and (ii) within the acyltransferase encoding gene (KL106-D3 found in JIE4005, JIE4019, and JIE4020) (Fig. 3a). All insertions caused the interruption of open reading frames producing characteristic 4 bp duplications (Fig. 3a). Sequences representative of the three new *cps* locus variants were deposited in GenBank. Capsule production was significantly different among the CP-*K. pneumoniae* strains (ANOVA, p < 0.001) (Fig. 3b). Production was significantly decreased in JIE2783 and JIE4282 (capsular variants KL106-D2 and KL106-D4 respectively), while it appeared significantly higher in JIE3095 (Fisher’s protected LSD, p < 0.05), though in this latter strain no sequence variation in the capsule encoding locus was identified.

**Figure 3.**
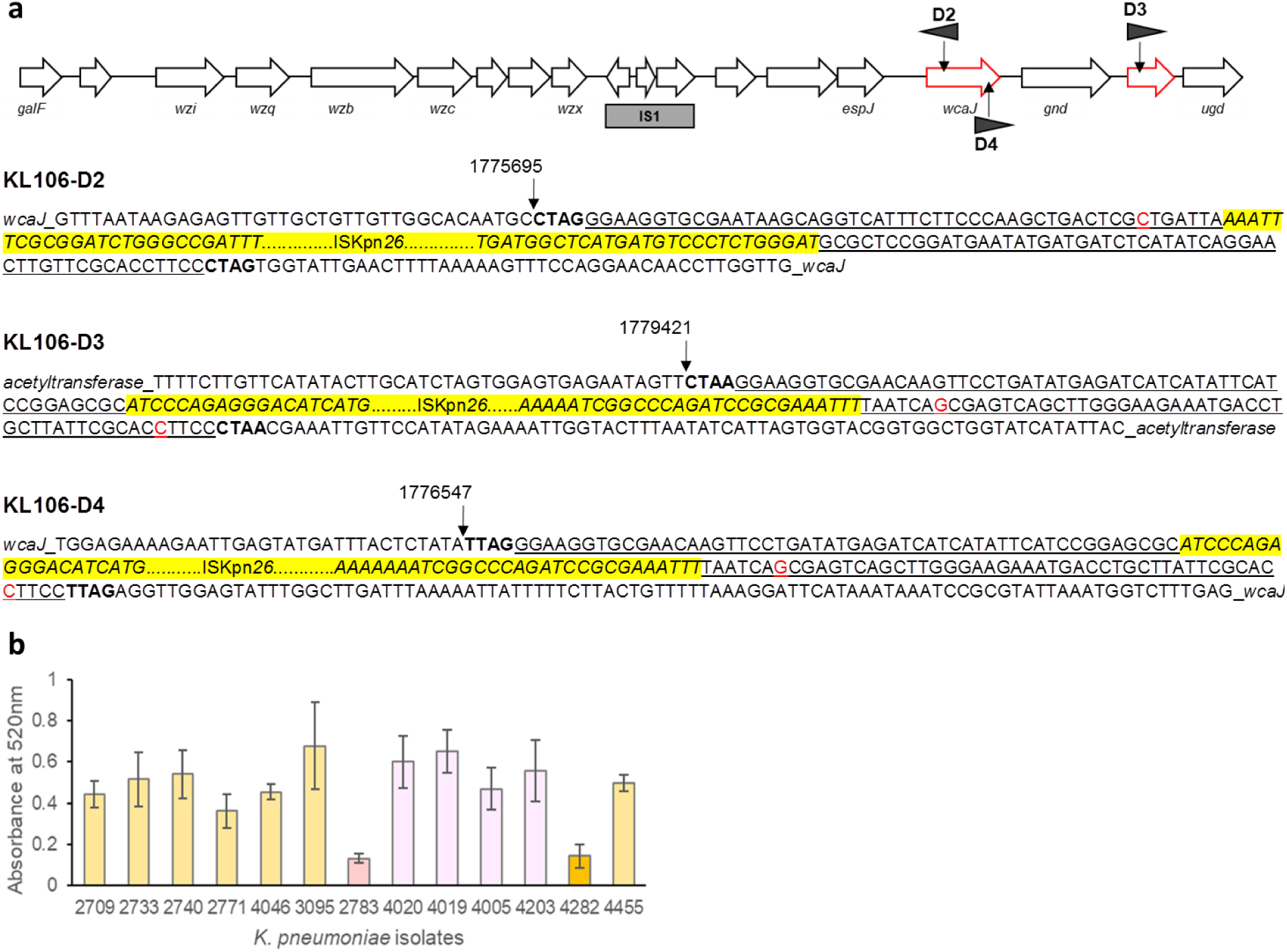
*cps* locus variants in *K. pneumoniae* ST258 clade 1 isolates. **a)** Schematic representing the *cps* locus gene arrangement in ST258 clade 1 isolates (KL106-D1; GenBank KR007677.1). Black triangles indicate ISKpn*26* insertions (directional) producing three variants (D2, D3 and D4). Drawing not to scale. Nucleotide sequence for ISKpn*26* insertions interrupting the *wcaJ* open reading frame (D2 and D4) and the acetyltransferase gene (D3) in the *cps* locus. In **bold**, insertion sequence (IS) direct repeats. <u>Underlined</u>, IS imperfect inverted repeats. ‘Yellow highlight’, indicates complete ISKnp*26* coding region. Nucleotide sequences of *cps* loci for variants KL106-D2 (JIE2783), KL106-D3 (JIE4005) and KL106-D4 (JIE4282) were deposited in GenBank. **b)** Capsule production assay. Capsule produced by ST258 clade 1 isolates was quantified based on uronic acid content (19,20). Results presented are average counts of three separate replicates.

In our set of strains, an evident association was observed between bacteriophage susceptibility profiles and capsular locus variants (Fig. 2) with very few of the tested phage (4/65; ∼ 6%) showing any cross-clade specificity. This correlation between capsule type and phage susceptibility held when a clade 1-specific (KL106) (AmPh_EK52), a clade 2-specific (KL107) (AmPh_EK38) and a phage with inter-clade range (JIPh_Kp122) were blind-tested on a larger panel (n=48) of CP-*K. pneumoniae* isolates from Europe (S3 Table).

#### Other cell surface structures

In contrast to the variable *cps* locus structure, gene content and arrangement in the LPS encoding loci was well conserved in our isolates. Accordingly, lipopolysaccharide profiles from silver staining showed a high degree of homogeneity with no significant differences in the O antigens of the short, long or intermediate chains (S1 Fig.). The lipid A component of JIE4282 differed in size from all other *K. pneumoniae* CG258 (S1 Fig.). According to Kleborate (26), JIE4282 is missing the *wbbM* gene encoding a glycosyltransferase required for *d*-galactan I biosynthesis (43). Of note, although JIE2793 (ST512) is missing one of the hypothetical proteins (*glmA*) in the O antigen operon (O2v2 type), this had no observable impact on the observed LPS profile.

Biofilm production levels also differed (unbalanced ANOVA, p < 0.001) and were found to be significantly higher in JIE2793, JIE3095 and JIE4282 compared to all others (Fisher’s protected LSD, p < 0.05) (S2 Fig.). Variable levels could be attributable to a number of factors, including variations associated with fimbrial genes. For instance, the *ecp* fimbrial operon associated with ROD-6 is missing in JIE4005, JIE4019, and JIE4020 (all KL106-D3) and in JIE4046 (S2 Table). JIE3095 also harbours a large 124 kb recombinant region which encompasses some fimbrial proteins. Limited capsule production (*e.g.* JIE4282) and expression of O-antigen variants (*e.g.* JIE2793) could also affect adhesion necessary for biofilm establishment. These variations in cell surface structures could be implicated with bacteriophage susceptibility, but no immediate correlation with phage host range was actually observed.

#### Prophages

The prophage content of clade 1 isolates was also variable. For example, a common prophage found in all other clade 1 representatives (designated phage 1 in S2 Table, with homology to viruses of the *Myoviridae* family) was truncated in JIE3095, missing most of the tail assembly module (Fig. 2). In JIE4005, 4019, and 4020 an additional unique prophage sequence was identified (Fig. 2; S2 Table). Based on prophage profile, clade 1 isolates could be subdivided into groups with broad association with bacteriophage susceptibility (Fig. 2).

#### Plasmids

All *K. pneumoniae* were MDR, and all ST258 isolates with the exception of JIE2740 carried the *bla*_KPC_ gene (Table 2; S4 Table). The overall plasmid signature for each ST258 clade was unique and remarkably uniform (Table 2; S3 Fig.). As expected (8,16), the *bla*_KPC_ allele 2 (*bla*_KPC-2_) was exclusively associated with ST258 clade 1, whilst *bla*_KPC-3_ was associated with clade 2 isolates and the closely related ST512 strain (Table 2). These genes were found exclusively on large (>20 kb) plasmids and co-localized by Southern hybridization with the IncFIIK *rep* gene (Table 2; S3 Fig.). All isolates carried multiple genes conferring extended-spectrum β-lactam resistance (ESBL) other than *bla*_KPC_ (S4 Table). Antibiotic resistance genotypes determined by WGS analysis accounted for all resistance phenotypes determined by standard clinical screening (S4 Table).

**Table 2.** Plasmid content in sequenced *K. pneumoniae* CG258 isolates.*

| Isolate<sup>^</sup> | ST258 clade | *bla<sub>KPC</sub>* allele | Plasmid replicons<sup>†</sup> | Plasmid sizes (kb)<sup>**</sup> |  |
| --- | --- | --- | --- | --- | --- |
|  |  |  |  | 20-100 | >100 |
| 2487 | na (ST11) | none | IncFIIK IncF-like ColE-like | nd | nd |
| 2713 | na (ST11) | none | IncFIIK ColE-like | nd | nd |
| **2709** | 1 | 2 | IncFIIK IncFIB-pKpQil-like IncX3 ColE-like | none | <u>242.5; 104.5</u> |
| 2733 | 1 | 2 | IncFIIK IncX3 ColE-like | **43** | <u>194</u> |
| **2740** | 1 | none | IncFIIK IncX3 ColE-like | 41 | <u>194</u> |
| 2771 | 1 | 2 | IncFIIK IncFIB-pKpQil-like IncX3 ColE-like | 41 | <u>104.5</u> |
| 2783 | 1 | 2 | IncFIIK IncFIB-pKpQil-like IncX3 ColE-like | **43** | <u>194; 104.5</u> |
| 3095 | 1 | 2 | IncFIIK IncX3 ColE-like | 41 | <u>165</u> |
| **4005** | 1 | 2 | IncFIIK IncX3 ColE-like | 43 | <u>160.5</u> |
| **4019** | 1 | 2 | IncFIIK IncF IncX3 ColE-like | 43 | <u>160.5</u>; 150 |
| 4020 | 1 | 2 | IncFIIK IncX3 ColE-like | 43 | <u>160.5</u> |
| 4046 | 1 | 2 | IncFIIK IncX3 ColE-like | 41 | <u>160.5</u> |
| 4203 | 1 | 2 | IncFIIK IncX3 ColE-like | 41 | <u>194; 104.5</u> |
| 4282 | 1 | 2 | IncFIIK IncFIB-pKpQil-like IncX3 ColE-like | 41 | <u>200; 110</u> |
| 4455 | 1 | 2 | IncFIIK IncFIB-pKpQil-like IncX3 ColE-like | 43 | <u>160.5</u> |
| 2793 | na (ST512) | 3 | IncFIIK IncN ColE-like | 58 | **208** |

Table 2 continued
| Isolate<sup>^</sup> | ST258 clade | *bla*<sub>KPC</sub> allele | Plasmid replicons<sup>†</sup> | Plasmid sizes (kb)<sup>**</sup> |  |
| --- | --- | --- | --- | --- | --- |
|  |  |  |  | 20-100 | >100 |
| 4626 | 2 | 3 | IncFIIK IncR IncX3 ColE-like | **43** | <u>208</u>; 121 |
| 4660 | 2 | 3 | IncFIIK IncR IncX3 ColE-like | **43** | <u>208</u>; 121 |

\*Data obtained from WGS analysis except where otherwise specified. <sup>^</sup>, in bold isolates from same patient; <sup>†</sup>, based on PlasmidFinder scheme implemented in BAP (30); \*\*, approximate sizes determined by S1-PFGE; nd, not determined. In bold, plasmids co-localizing with the *bla*<sub>KPC</sub> gene as identified by Southern blot hybridization. Underline, IncFIIK replicons detected in Southern blot hybridization.

### Bacteriophage characterization

Among the tested phages, we identified five unique double-stranded DNA bacteriophages AmPh_EK29, AmPh_EK52, AmPh_EK80, JIPh_Kp122, and JIPh_Kp127 that selectively target CP-*K. pneumoniae* ST258 clade 1 isolates, some of which may have therapeutic potential (Table 3; Table 4). WGS of purified viral DNA produced 11,340 to 5,049,732 reads that *de novo* assembled into one contig in all instances (S1 Table). Bacteriophage genomes size varied between 40.7 and 169.3 kb and GC content was lower than 50% (i.e. host genome) in all except for AmPh_EK52 (GC% 52.9) (Table 4; Fig. 4). No lysogeny or virulence associated genes were identified in these bacteriophage sequences, consistent with their lytic nature and thus indicating their suitability for therapeutic use (Fig. 4). The high degree of sequence similarity (>95%) to characterized *K. pneumoniae*-specific phage in the NCBI database and TEM imaging indicated that each selected phage belonged to the order Caudovirales (Table 4; Fig. 4).

**Table 3.** Efficiency of plating of selected CP-K. pneumoniae ST258 clade 1 bacteriophages^*^.

| Isolate (JIE) | ST | Clade | AmPh_EK29 |  | AmPh_EK52 |  | AmPh_EK80 |  | JIPh_Kp122 |  | JIPh_Kp127 |  |
| --- | --- | --- | --- | --- | --- | --- | --- | --- | --- | --- | --- | --- |
| 2713 | ST11 | NA | none | 4 | none | 4 | none | 4 | *** | 3 | *** | 3 |
| 2487 | ST11 | 1 | none | 4 | none | 4 | none | 4 | *** | 3 | *** | 3 |
| 4203 | ST258 | 1 | none | 4 | 1.0x10<sup>9</sup> (125) | 1 | none | 4 | *** | 3 | *** | 3 |
| 4046 | ST258 | 1 | 1.1x10<sup>9</sup> (55) | 1 | 8.0x10<sup>8</sup> (100) | 1 | *** | 3 | *** | 3 | *** | 3 |
| 4020 | ST258 | 1 | none | 4 | 4.0x10<sup>9</sup> (500) | 1 | *** | 3 | *** | 3 | *** | 3 |
| 4019 | ST258 | 1 | none | 4 | 2.6x10<sup>9</sup> (325) | 1 | *** | 3 | *** | 3 | *** | 3 |
| 4005 | ST258 | 1 | none | 4 | 1.6x10<sup>9</sup> (200) | 1 | *** | 3 | *** | 3 | *** | 3 |
| 3095 | ST258 | 1 | 4.6x10<sup>8</sup> (23) | 1 | 8.0x10<sup>8</sup> (100) | 1 | *** | 3 | *** | 3 | *** | 3 |
| 2709 | ST258 | 1 | **2.0x10<sup>9</sup> (100)** | **1** | **8.0x10<sup>8</sup> (100)** | **1** | 4.0x10<sup>8</sup> (15) | 1 | *** | 3 | *** | 3 |
| 2733 | ST258 | 1 | 8.0x10<sup>8</sup> (40) | 1 | 6.0x10<sup>8</sup> (75) | 1 | 4.0x10<sup>8</sup> (15) | 1 | *** | 3 | *** | 3 |
| 2740 | ST258 | 1 | 4.0x10<sup>8</sup> (20) | 1 | 2.6x10<sup>9</sup> (325) | 1 | none | 4 | *** | 3 | *** | 3 |
| 2771 | ST258 | 1 | 1.8x10<sup>9</sup> (90) | 1 | 4.0x10<sup>8</sup> (50) | 1 | *** | 3 | *** | 3 | *** | 3 |
| 2783 | ST258 | 1 | 2.4x10<sup>9</sup> (120) | 1 | none | 4 | 6.0x10<sup>9</sup> (100) | 1 | 2.5x10<sup>9</sup> (313) | 1 | 1.5x10<sup>7</sup> (188) | 1 |
| 4455 | ST258 | 1 | 2.0x10<sup>9</sup> (100) | 1 | 2.5x10<sup>8</sup> (31) | 1 | *** | 3 | *** | 3 | *** | 3 |
| 4282 | ST258 | 1 | none | 4 | none | 4 | **6.0x10<sup>9</sup> (100)** | **1** | **8.0x10<sup>8</sup> (100)** | **1** | **8.0x10<sup>6</sup> (100)** | **1** |
| 2793 | ST512 | NA | 6.0x10<sup>8</sup> (30) | 1 | none | 4 | none | 4 | *** | 3 | *** | 3 |
| 4660 | ST258 | 2 | none | 4 | none | 4 | none | 4 | *** | 3 | *** | 3 |
| 4626 | ST258 | 2 | none | 4 | none | 4 | none | 4 | *** | 3 | *** | 3 |

\*, in brackets percentage EOP when compared to amplification strain. \*\*\*, indicates clearing in bacterial lawn without single plaque formation (confluent lysis or abortive lysis). Bold, original amplification host. Scores indicate: 1, high titre lysis; 3 and 4, poor or no lysis.

**Table 4.**
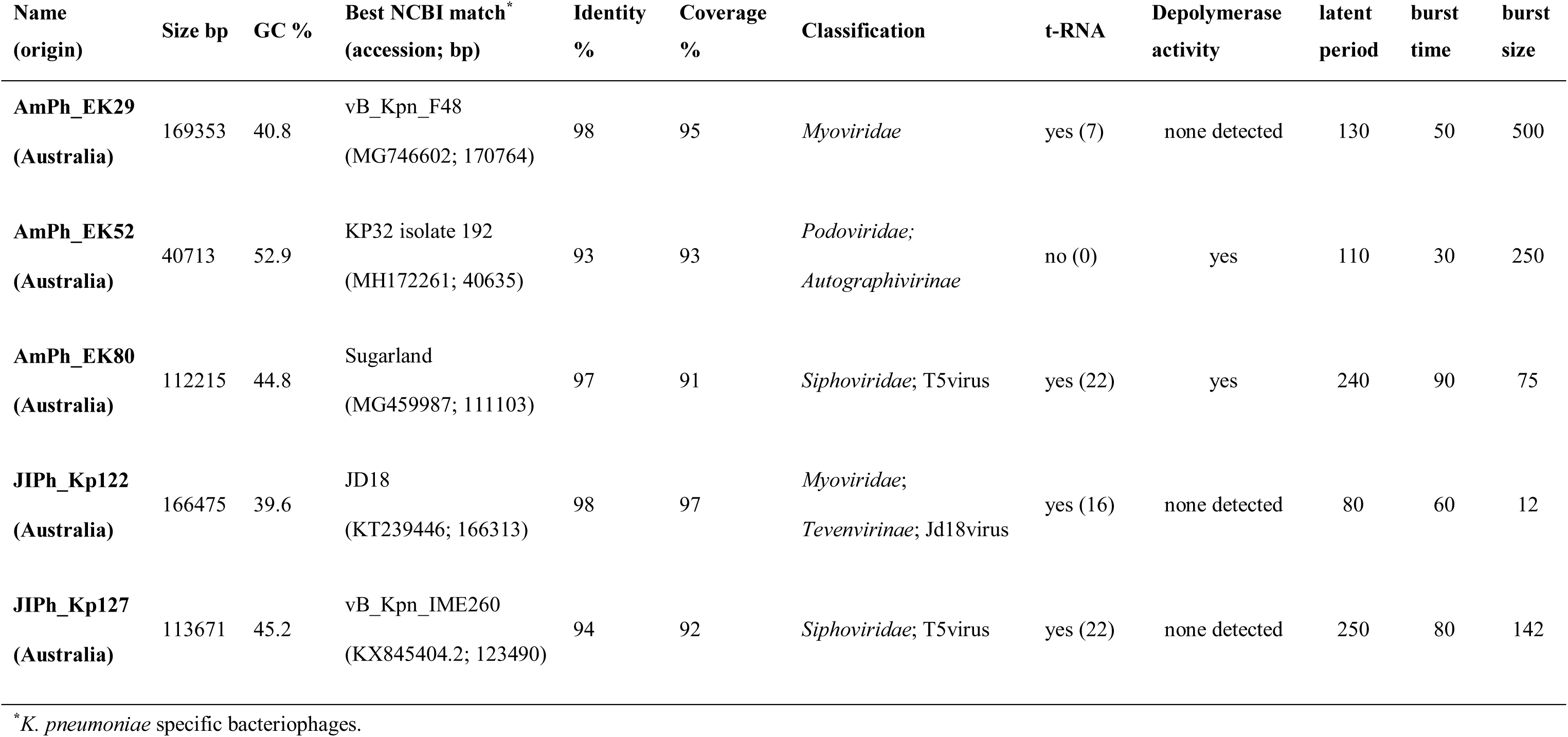
Characteristics of selected *K. pneumoniae* ST258 clade 1-specific bacteriophages.

| Name<br>(origin) | Size bp | GC % | Best NCBI match*<br>(accession; bp) | Identity<br>% | Coverage<br>% | Classification | t-RNA | Depolymerase<br>activity | latent<br>period | burst<br>time | burst<br>size |
| --- | --- | --- | --- | --- | --- | --- | --- | --- | --- | --- | --- |
| **AmPh_EK29**<br>(Australia) | 169353 | 40.8 | vB_Kpn_F48<br>(MG746602; 170764) | 98 | 95 | *Myoviridae* | yes (7) | none detected | 130 | 50 | 500 |
| **AmPh_EK52**<br>(Australia) | 40713 | 52.9 | KP32 isolate 192<br>(MH172261; 40635) | 93 | 93 | *Podoviridae*;<br>*Autographivirinae* | no (0) | yes | 110 | 30 | 250 |
| **AmPh_EK80**<br>(Australia) | 112215 | 44.8 | Sugarland<br>(MG459987; 111103) | 97 | 91 | *Siphoviridae*; T5virus | yes (22) | yes | 240 | 90 | 75 |
| **JIPh_Kp122**<br>(Australia) | 166475 | 39.6 | JD18<br>(KT239446; 166313) | 98 | 97 | *Myoviridae*;<br>*Tevenvirinae*; Jd18virus | yes (16) | none detected | 80 | 60 | 12 |
| **JIPh_Kp127**<br>(Australia) | 113671 | 45.2 | vB_Kpn_IME260<br>(KX845404.2; 123490) | 94 | 92 | *Siphoviridae*; T5virus | yes (22) | none detected | 250 | 80 | 142 |

\**K. pneumoniae* specific bacteriophages.

**Figure 4.**
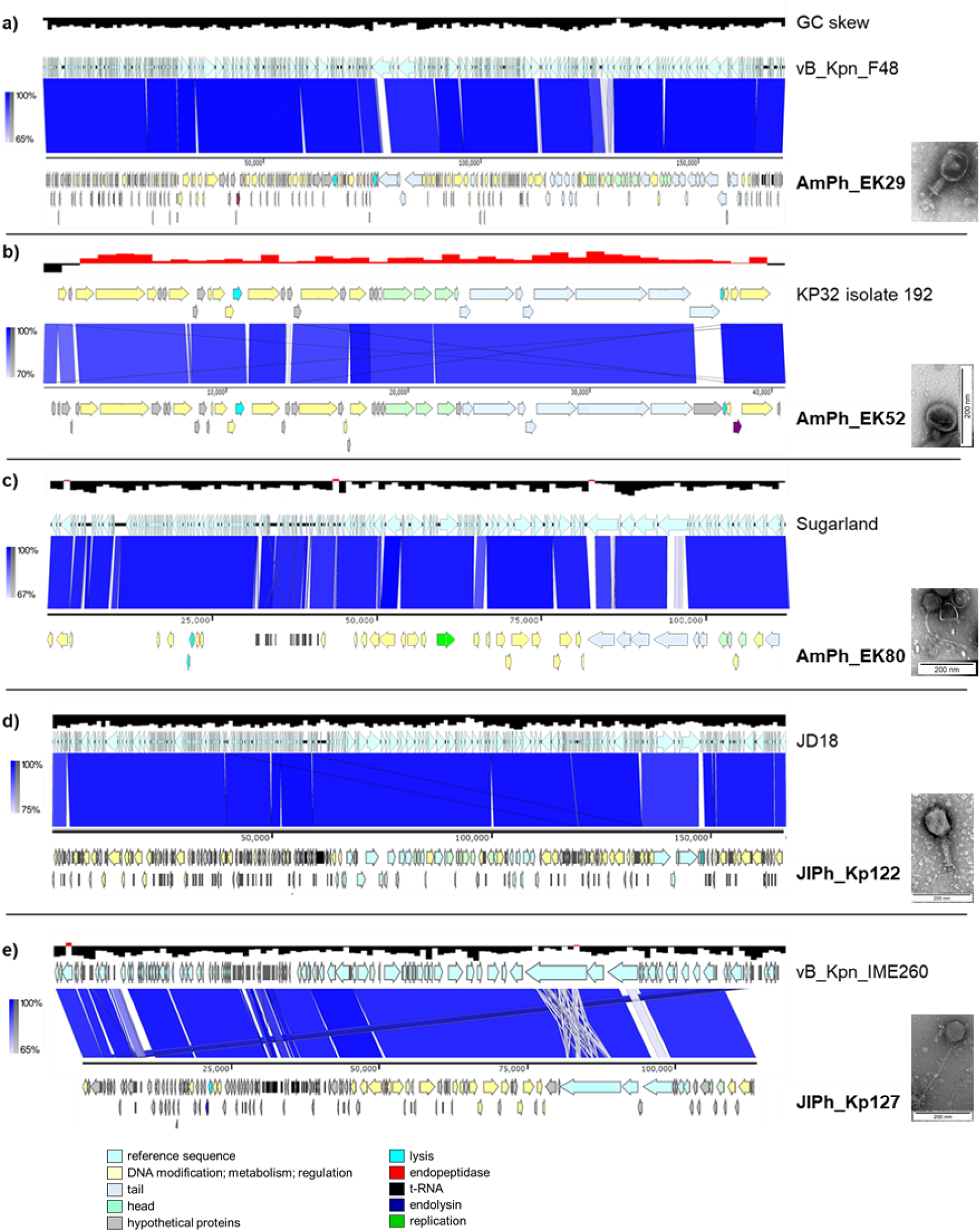
Comparative genome analysis of five sequenced *K. pneumoniae* ST258 bacteriophages. Schematics were produced using EasyFig (32) and show the structural organization of the selected phages compared to their best match (>95% nucleotide homology) reference in the GenBank database. Genes are color coded according to function. Phage morphology was captured using transmission electron microscopy.

AmPh_EK29 and JIPh_Kp122 (*Myoviridae*-like), presented a prolate head (approx. 80 by 100 nm) and contractile tale (100 nm) (Table 4; Fig. 4a and 4d). AmPh_EK52 resembled *Podoviridae* phages for having a small thick non-contractile tail (approx. 20 nm long) and close homology to other members of this family including genome size of about 40 kb and absence of t-RNAs in its genome (Table 4; Fig. 4b). JIPh_Kp127 and AmPh_EK80 were both T5-like *Siphoviridae* viruses with long thin non-contractile tails (Table 4; Fig. 4c and 4e). Screening of entries in the NCBI database by BLASTn identified close relatives of these bacteriophages but no identical sequences (Table 4), and in genome comparisons with best matching GenBank entries the modular structure and order were preserved in all cases, with the main regions of difference found in tail or tail-associated open reading frames (Fig. 4).

All five bacteriophages efficiently lysed target bacteria *in vitro* at high titre and in combination captured the entire ST258 clade 1 subset (Table 3). However, host ranges were unique for each phage. AmPh_EK80, JIPh_Kp122 and JIPh_Kp127 showed limited activity toward ST258 strains, bar a few specific targets that lysed at high titre, and often were associated with confluent lysis zones, indicating the possibility of lysis from without or abortive lysis (Table 3). All phages were highly stable in SM buffer maintaining high titre at a range of temperatures (4°C for >1 month; 21°C for one week; 37°C for 24 h) and pH levels (pH 3, 6, 7 and 8 for 4 h). Exposure to chloroform at 4°C decreased the stability of AmPh_EK29 by 2-3 orders of magnitude, but had no effect on the stability of the remaining four phages. One-step growth curves revealed latent periods of 80-250 min and burst sizes between 12 and 500 pfu/cell (Table 4; S4 Fig.). Growth curves for AmPh_EK80 and JIPh_Kp127 were comparable. Phage JIPh_Kp122 had the shortest latent time (80 min), while AmPh_EK52 had the shortest burst time (30 min).

#### Accession number(s)

The Illumina sequencing datasets of all *K. pneumoniae* isolates obtained in this work were deposited in SRA (NCBI) database under Bioproject PRJNA529495. The *cps* locus variants were deposited in the GenBank database (NCBI) under accessions: XXXXX for KL106-D2 (JIE2783), XXXXX for KL106-D3 (JIE4019), and XXXXX for KL106-D4 (JIE4282). The complete annotated genomes of *K. pneumoniae* phages AmPh_EK29, AmPh_EK52, AmPh_EK80, JIPh_Kp122, and JIPh_Kp127 were also deposited in the GenBank database (NCBI) under accession numbers XXXXX, XXXXX, XXXXX, XXXXX and XXXXX, respectively.

## Discussion

The increasing challenges posed by the rise of antibiotic resistance in human pathogens have revitalized interest in the use of bacteriophage for the treatment of bacterial infections (10-12). Among MDR pathogens, CP-*K. pneumoniae* is a serious clinical concern, as both gut colonizer and agent of severe sepsis when invading sterile body sites (1,44). In this study, ST258 isolates were predominant in the local clinical CP-*K. pneumoniae* population, with overrepresentation of ST258 clade 1 (KL106-D1), reflecting the epidemiology of a recent local outbreak (9). The incidence of *bla*_KPC_ in Australia has been rather limited when compared to its dissemination in other countries (16). The Western Sydney Health district is one of the largest in the country and we could only identify 0.2% CP-*K. pneumoniae* with *bla*_KPC_ over the past six years in its diagnostic enterobacterial collection (JIE), which is in line with reported national frequencies (45). However, as our own data confirms, tracking of these pathogens remains paramount due to the consistent association of multidrug resistance with mobilizable elements (16,36).

In the *K. pneumoniae* genome, the *cps* locus, encoding the capsular polysaccharide outer layer, is a recognized recombination hotspot, responsible for the diversification of clonal lineages, particularly within the CG258 group (6-8), and over 100 capsular types have been identified in this species (42,46). The capsule is a complex structure of repeating sugar subunits that protects the cell from external threats (including phage attack) and enhances *K. pneumoniae* virulence, being implicated in resistance to host defence mechanisms, immune evasion, adherence, and biofilm formation (47). *K. pneumoniae* types produce capsule in varying degrees and hypermucoviscosity (excessive capsule production) has been linked to increased virulence (47). In our study, we identified three novel capsular variants in ST258 clade 1 due to ISKpn*26* insertion in the *cps* locus, and responsible for the intra-clade variation in our population which correlated remarkably well with reduced host range for many of the clade 1 infecting phages. ISKpn*26* has been recently implicated in unique recombination events in MDR *K. pneumoniae* (48) and may be worthwhile tracking. Two of the variants (D2 and D4), where the IS interrupted the *wcaJ* gene, were found in isolates with reduced capsule production in accordance with studies demonstrating that disruption of this gene has a detrimental effect on capsule content (46,49). The positive correlation between phage resistance and reduced virulence has been previously observed in the adaptive interplay between *Klebsiella* and its bacteriophages, with the isolation of less virulent bacterial mutants, and may be an expected trade-off in their evolutionary arms race (50). It is therefore not surprising that co-evolution of specific phage with this host resulted in the narrow host ranges observed in this study, a characteristic that could be exploited for therapeutic purposes, as specific recognition by phage receptors (tail spikes) of complementary anti-receptors on the host cell surface is at the core of phage lytic activity (51,52).

Comparative analysis of genomic data and bacteriophage susceptibility profiles showed a marked association between the host range of most of the tested viruses with capsular type. Very few examples of cross-reactivity were observed among *K. pneumoniae* CG258 phages that lysed different capsular types with high specificity. Host range was further restricted within clades, i.e. no phage lysing ST258 clade 1 effectively also infected clade 2 isolates, and *vice versa*. Phage AmPh_EK29 as a notable exception lysed only a subset of ST258 clade 1 strains (capsular type KL106), but also the ST512 representative isolate (capsular type KL107), indicative of a unique receptor specificity for this virus. JIPh_Kp122 and JIPh_Kp127 also showed broader spectra beyond capsular type, but lysed ST258 isolates overall with poor efficiency, with evidence of abortive or passive lysis (53), perhaps limiting their therapeutic value. Capsule-targeting bacteriophages have been shown to be effective against *K. pneumoniae* of different capsular type *in vitro* and *in vivo*, along with depolymerases, which are enzymes produced by phages that cleave glycosidic bonds disrupting capsule integrity (54,55). Depolymerase activity was originally shown to be random, however recent work demonstrated depolymerase specificity toward certain K types (56,57), an indication that enzymatic degradation may be causing the patterns of lytic activity observed in our study. We did not find immediate evidence that all our phage encoded these proteins and, for the ones for which a depolymerase-related phenotype was observed, we could not readily identify the putative coding regions, except for AmPh_EK52.

Even though the attachment mechanisms or receptor binding sites for any of the tested phages were not investigated, host range matching with genomic data allowed for the selection of a number of viruses with unique characteristics that could be further examined for therapeutic applications. All the phages sequenced in this work were highly homologous to previously characterized lytic viruses shown to be effective against specific *K. pneumoniae* STs. However, none matched the specificities (host ranges) against ST258 of our viruses. Comparative analysis of phage genomes identified preferential loci of variability, mostly related with tail fibres or tail-associated genes, likely the primary receptors responsible for each specific host range. The different but complementary host ranges of the phages characterized here may indicate differences in receptor specificity (see AmPh_EK52 versus JIPh_Kp122 for example) and could lead to successful therapeutic combinations (high activity, poor resistance development).

The complex dynamic interactions of phages and their hosts and the co-evolution mechanisms at play have undermined the direct prediction of phage susceptibility from host genomics, even when dealing with clonal populations. Though capsular variation was at the core of the divergence of isolates within this subset, fine genomic diversity in our ST258 clade 1 strains was also associated with other elements such as prophage content, porin defects, and plasmid content. The ST258 clade 1 subset in this study roughly divided into three subgroups based on prophage profile, further evidence of the genome plasticity in these species (58). Prophage content was likely implicated in resistance to some of the tested phages (e.g. AmPh_EK29) and must also be considered when designing optimal therapeutic mixes. Variability in the lipid A core of the LPS surface layer could also contribute phage resistance in JIE4282, as this is generally highly conserved within *K. pneumoniae*, making it an important determinant for host receptor recognition (51,52). Other differences (lack of fimbrial locus in JIE4046, different plasmid content (JIE4005, 4019 and 4020), *ompK36* variants etc.) seemed not to directly impact the susceptibility patterns to most of the tested phages. However, these all have the potential to affect viral host range and synergy when phages are to be used for therapeutic cocktails preparation (51,52).

The unique relationship between *K. pneumoniae* and its phages highlighted in this study is very important for future progress in therapeutic applications specifically targeting ST258 isolates or other hypervirulent types Better bioinformatic tools and larger well characterized microbial collections may allow for better definition of predictive algorithms. As we have demonstrated here capsular variation is critical to phage susceptibility and must be considered when designing effective therapeutics against this pathogen.

## Acknowledgements

The authors would like to acknowledge Alicia Arnott, Nathan Bachmann, Chayanika Biswas, Rajat Dhakal, Elena Martinez, Ranjeeta Menon, Rebecca Rockett, Rosemarie Sadsad, Verlaine Timms, Qinning Wang and Vitali Sintchenko at the Pathogen Genomics Unit, Centre for Infectious Disease and Microbiology –Public Health, Westmead Hospital, for their assistance with whole genome sequencing; Dongwei Wang for her assistance with TEM; Sydney Water, Murray McDermott and Graziano (Rowland Village), and Lee Thomas for their help with specimen collection for bacteriophage isolation.

